# NinjaSeq: programmable restriction enzyme-based sequencing library preparation with random access for DNA data storage

**DOI:** 10.64898/2026.06.11.730843

**Authors:** Ignas Galminas, Omer Sabary, Hadas Abraham, Kornelija Kaminskaitė, Tomer Cohen, Vakarė Gruodytė, Gediminas Alzbutas, Zohar Yakhini, Rūta Palepšienė, Lukas Žemaitis, Eitan Yaakobi, Simonas Juzenas

**Affiliations:** Faculty of Natural Sciences, Vytautas Magnus University, Kaunas, Lithuania; Department of DNA data storage, Genomika, Kaunas, Lithuania; Computer Science, Technion-Israel Institute of Technology, Haifa, Israel; Computer Science, Reichman University, Herzliya, Israel; Institute of Biotechnology, Life Sciences Center, Vilnius University, Vilnius, Lithuania

**Keywords:** DNA data storage, REases, nanopore sequencing, random access

## Abstract

DNA data storage allows sequences to be defined without biological constraints, yet readout workflows still depend on generic end-repair/dA-tailing chemistry. We developed NinjaSeq, a type IIS restriction endonuclease library-preparation strategy that incorporates recognition sites into primer flanks, enabling digestion to generate adapter-compatible overhangs and eliminating the need for conventional end preparation. By combining this chemistry with constrained coding that excludes internal recognition motifs, NinjaSeq produced sequencing quality and decoding performance consistent with standard protocols while reducing reagent burden and simplifying processing, including compatibility with one-pot restriction-ligation. The same sequence-directed design also enables physical random access during library preparation: targeting file-specific flanking sites enriched a desired file from a mixed pool by about sixteen-fold in a proof-of-concept experiment. These results position NinjaSeq as a practical ONT readout approach for DNA data storage.

**HIGHLIGHTS:**

- NinjaSeq replaces end-repair/dA-tailing with REases for nanopore sequencing
- Constrained encoding excludes recognition motifs to protect payloads from cleavage
- NinjaSeq achieves decoding accuracy comparable to standard library preparation
- Designing file-specific RRS enables random access during library preparation

## INTRODUCTION

The exponential expansion of digital information is rapidly surpassing the capabilities of conventional storage technologies, motivating the search for alternative media with superior density, longevity, and reliability. DNA has emerged as a particularly promising solution, offering theoretical storage densities orders of magnitude higher than existing systems, remarkable chemical stability under appropriate conditions^1–3^, and mature biochemical techniques for its accurate replication and sequencing^4^. These features together ensure that DNA can support both reliable duplication and retrieval of information. The practical deployment of DNA-based storage is further facilitated by advances in sequencing technologies, which have greatly improved the efficiency and scalability of information retrieval. Among these, Oxford Nanopore Technologies (ONT) sequencing enables real-time, portable, and high-throughput reading of DNA, providing a flexible platform for rapid and scalable decoding of DNA-encoded data^5^.

ONT library preparation for nanopore sequencing typically comprises two steps. First, end preparation generates DNA fragments bearing 5′-phosphate groups and 3′ adenylated (A) overhangs. This is commonly achieved using an end-repair/dA-tailing enzyme mix (DNA polymerase activity) together with a kinase activity for 5′ phosphorylation (e.g., T4 polynucleotide kinase)^6^. Second, the adenylated fragments are ligated to sequencing adapters carrying complementary 3′-thymine (T) overhangs, typically using T4 DNA ligase. Adapter ligation is essential because ONT adapters include a 5′ motor protein that promotes strand unzipping and regulates single-stranded DNA (ssDNA) translocation through the nanopore. As ssDNA passes through the pore, characteristic current disruptions are recorded and subsequently converted into nucleotide sequences^5^.

Despite its effectiveness, the standard workflow is multi-step and relies on multiple enzyme mix, increasing cost and operational complexity. This motivates simplified, more cost-effective protocols based on easily available enzymes. In DNA-based data storage, such simplification is especially feasible because the encoded sequences are designed in silico and can be constrained to satisfy additional biochemical requirements. Specifically, restriction enzyme recognition sequences (RRSs) can be reserved for the flanking primer regions and excluded from the payload, thereby preventing unintended internal cleavage within information-bearing DNA.

NinjaSeq builds on this premise by using restriction endonucleases (REases), which cleave DNA at defined recognition sequences with high efficiency and specificity and leave 5′-phosphate termini after cleavage. By selecting enzymes whose recognition sites generate the required adenine-containing overhangs, DNA fragments can be made directly compatible with ONT adapter ligation without a separate end-preparation step. In NinjaSeq, the target DNA is first amplified by PCR using primers that introduce an REase-specific RRS at both termini; the selected REase then cleaves at these sites to produce adapter-compatible overhangs, and the resulting fragments are ligated to ONT adapters using T4 DNA ligase to generate sequencing-ready libraries (**Fig. 1a**).

**Figure 1.**
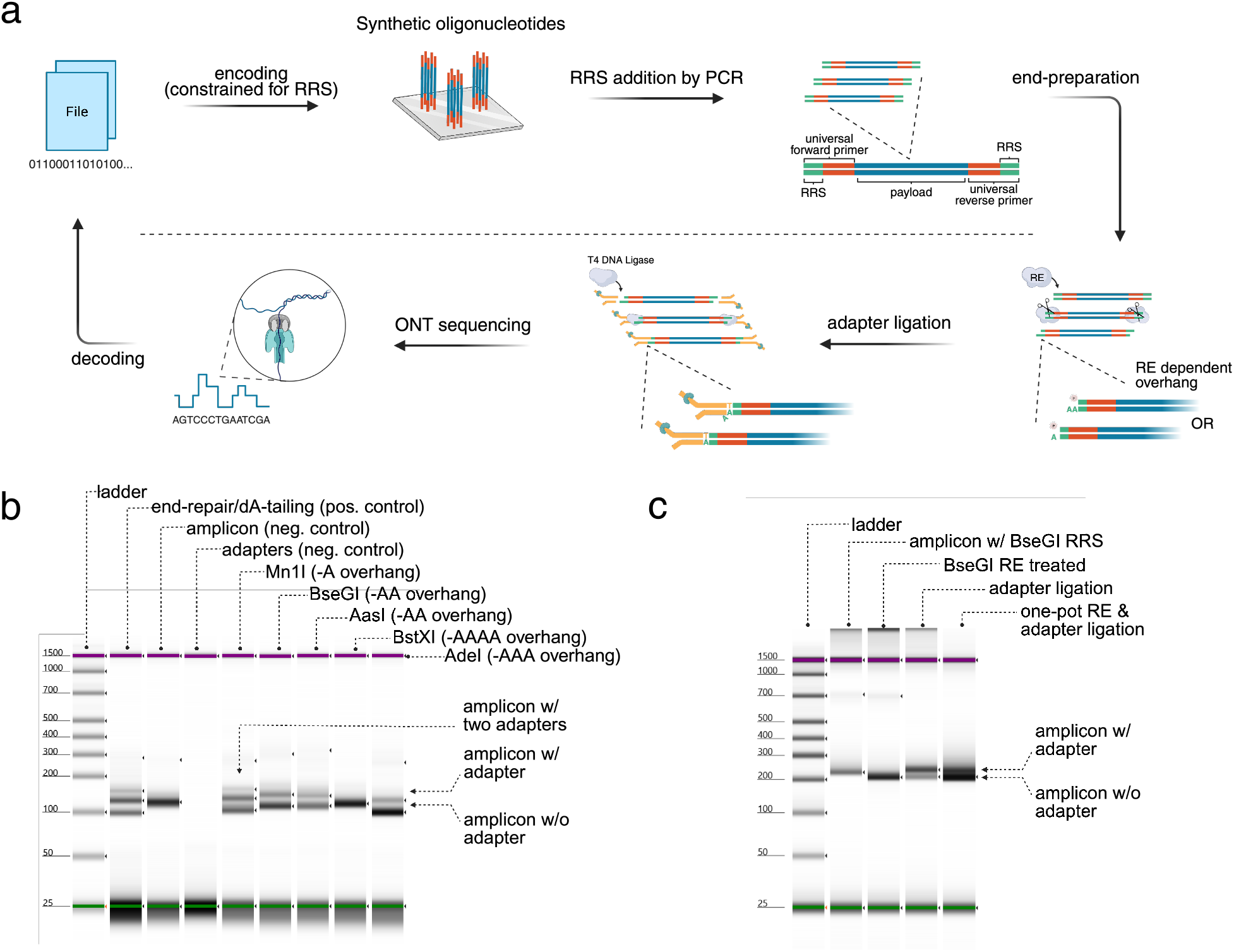
Concept of restriction endonuclease-based sequencing library preparation for DNA-based data storage applications. **a)** Schematic workflow of restriction endonuclease-based ONT nanopore sequencing library preparation for DNA data storage applications. **b)** Proof of concept for adapter ligation to an oligopool using multiple restriction enzymes that leave single or multiple adenine (A) 3’ overhangs and T4 DNA ligase. The figure displays electrophoresis results showing adapter ligation efficiency. **c)** Verification experiment for restriction endonuclease-based adapter ligation. Electrophoresis results demonstrate successful adapter ligation using BseGI restriction endonuclease and T4 DNA ligase, carried out as separate reactions or as a one-pot reaction. Created with BioRender.com.

The NinjaSeq workflow offers three key advantages over standard library preparation protocols. First, by utilizing widely available restriction enzymes rather than specialized end-repair/dA-tailing kits, it substantially reduces reagent costs. Second, the compatibility of the enzymatic steps allows for a “one-pot” reaction, simplifying the workflow and minimizing sample handling. Finally, the sequence-specific nature of REases enables inherent random access capabilities during library preparation, as specific data files can be selectively processed by targeting unique RRSs embedded in their primers.

## RESULTS

### Concept of restriction endonuclease-based sequencing library preparation (NinjaSeq)

To evaluate whether REases can mimic end-repair/dA-tailing, the enzymes were selected based on their ability to be programmable (contain Ns in RRS) and generate A-containing overhangs. In particular, the selection included enzymes that produce different overhang lengths, including -A (MnlI), -AA (BseGI, AasI), -AAA (AdeI), and -AAAA (BstXI). For each enzyme, adapter ligation was performed and the resulting products were compared with those from a standard end-repair/dA-tailing reaction (**Fig. 1b**). Effective ligation was obtained when REase digestion generated 1–3 nt adenine overhangs (-A, -AA, or -AAA). In contrast, ligation was not effective when the cleavage produced a four-adenine overhang (-AAAA). Among the tested overhangs, shorter -A ends (MnlI) yielded the highest ligation efficiency and was comparable with the end-repair/dA-tailing reaction. REases generating longer 2–3 nt adenine overhangs tended to produce products consistent with predominantly single-adapter ligation (**Fig. 1b**). Notably, even with -AA overhangs, ligation preferentially occurred at the innermost -A immediately adjacent to the 5′-phosphate on the top strand (**Supplementary Fig. 1**).

Further, the REase-based “end-preparation” was evaluated for compatibility with a single-pot restriction–ligation reaction in which cleavage and adapter ligation proceed in the same reaction. Technically, this resembles Golden Gate assembly^7^, in which concurrent restriction digestion and ligation are used to couple end generation and joining within a single reaction mixture. Such coupling is generally not possible in standard end-repair/dA-tailing protocols because end repair/dA-tailing enzymes possess strong 3’ to 5’ exonuclease activity that would degrade the 3’ T-overhang of the sequencing adapters if they were present in the reaction mix simultaneously.

To test one-pot ligation, the BseGI (-AA) restriction enzyme was used together with T4 DNA ligase in a single reaction (one-pot), or in a sequential workflow in which digestion and ligation were performed separately, including a purification step in between (two-pot). Mock adapters lacking the motor protein were used to evaluate ligation efficiency in both conditions. The results showed that adapters could be ligated to the amplicon library in the one-pot reaction, although at a lower efficiency than in the two-pot workflow (**Fig. 1c**). This reduced efficiency may reflect competition from re-ligation of the short (~6 nt) fragments released by BseGI cleavage, which can re-join to the amplicon ends and thereby reduce the effective yield of adapter-ligated products.

Taken together, these results indicate that REase digestion can substitute for end-repair/dA-tailing when it generates short (1–3 nt) A-overhangs. Moreover, one-pot restriction–ligation is feasible, although less efficient than the two-pot workflow.

### Constrained data encoding for restriction endonuclease-based DNA data storage applications

In order to use REase-based end-preparation for payload-encoded libraries, it is desirable to ensure that restriction enzyme recognition sites (RRS) do not occur within the payload sequence, because internal sites would be cleaved during library preparation and compromise the construct. RRS avoidance can therefore be formulated as a constrained-encoding problem: given the DNA alphabet {*A, C, G, T*} and a set of forbidden substrings *F*, valid payloads are restricted to the constrained code *C*(*F*) (the set of all DNA sequences that avoid every motif in *F*). Avoiding these motifs should help to prevent unintended restriction cuts and improve library integrity. However, excluding motifs also removes many otherwise valid payload sequences, reducing how much information can be packed into a given sequence length. This loss can be quantified by the constrained-system capacity, defined as the maximum achievable encoding rate (bits per nucleotide) under the chosen motif-avoidance rules.

To analyze and implement these constraints, *C*(*F*) can be represented as a finite labelled directed constraint graph, in which each state tracks the relevant recent suffix context needed to ensure that extending the sequence does not introduce any forbidden motif. In this representation, paths through the graph correspond to valid sequences, enabling both efficient generation of allowed payloads and quantitative evaluation of the information-rate penalty via the graph’s adjacency structure.

As a concrete example, the constraint graph for avoiding the BseGI RRS motif GGATG is shown in **Fig. 2a**. The states (nodes) represent partial matches to the forbidden motif, {Φ, *G, GG, GGA, GGAT*}, and the transitions (edges) indicate which nucleotide can be added next without forming GGATG. This graph provides a compact finite-state description of all payload sequences that avoid the motif and serves as a representative example of how RRS avoidance constraints can be encoded and analyzed. Although not shown in this example, the reverse-complement of each RRS must also be excluded to fully prevent internal restriction sites on either strand.

**Figure 2.**
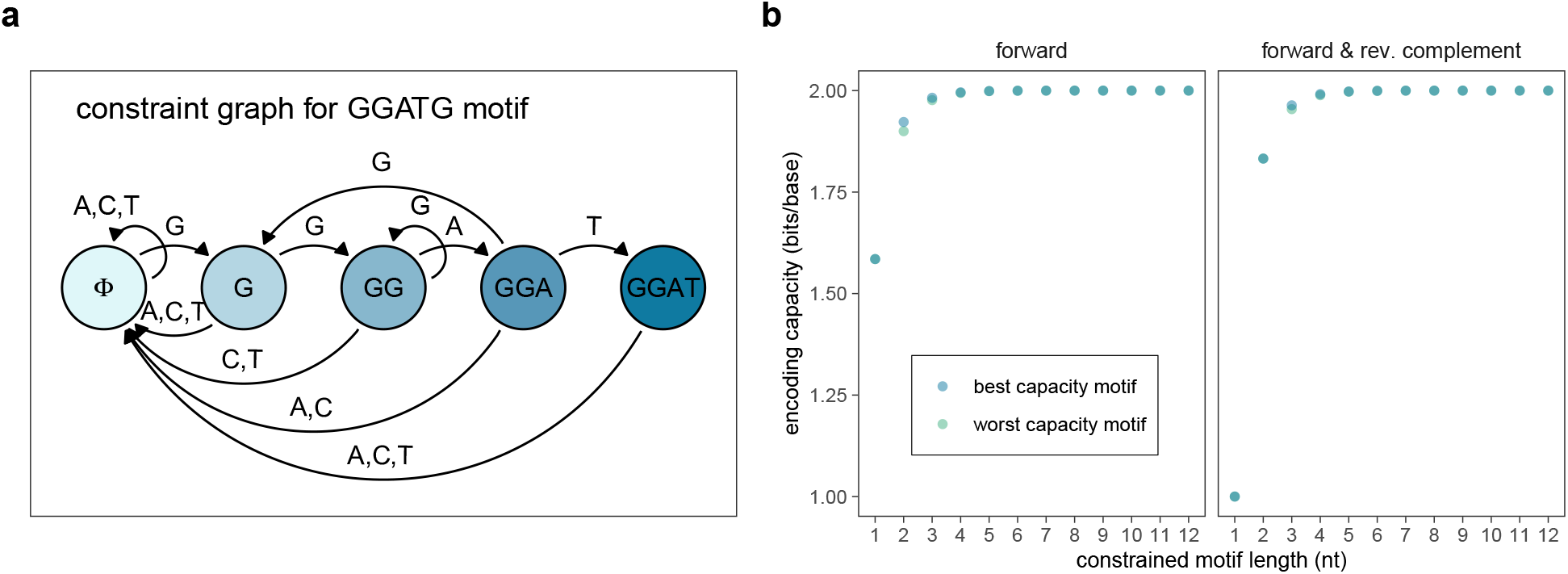
Constrained encoding for RRS avoidance and encoding capacity. **a)** Constraint graph for avoiding the BseGI recognition motif (GGATG). The nodes represent the state (corresponding to the length of the current suffix matching the forbidden motif), while directed edges indicate allowed nucleotide transitions that prevent the formation of the full restriction site. **b)** Theoretical encoding capacity of constrained systems as a function of the forbidden motif length (nt). The plot displays the maximum achievable information rate (bits per nucleotide) for the most restrictive (worst capacity; autocorrelating) and least restrictive (best capacity; homopolymer) motif compositions at each length.

To evaluate encoding capacity across practical REase choices, capacities were computed for a range of forbidden sets and analyzed as a function of recognition-site length and motif sequence composition. For each recognition-site length, capacities were determined for both the least and most restrictive motifs based on sequence composition: homopolymer motifs, which typically allow the highest capacity, and autocorrelating motifs, where a sequence’s prefix is also its suffix, which yield the lowest capacity^8^. The results indicate that the capacity penalty decreases rapidly as the forbidden motif length *n* increases (**Fig. 2b**). When avoiding motifs on both the forward and reverse complement strands, the constraint is more significant for very short motifs (*n* = 2), with capacity dropping to approximately 1.83 bits per nucleotide. However, for typical RRS lengths of 4–6 nucleotides, the capacity remains high, ranging from roughly 1.989 (*n* = 4) to nearly 1.999 (*n* = 6) bits per nucleotide, regardless of motif composition. This value represents the amount of information per symbol when encoding sequences that avoid the forbidden pattern. It is important to note that this capacity is asymptotic. For any target value below the capacity, it is possible to design an encoder that achieves that information per symbol.

### Sequencing performance of NinjaSeq libraries from RRS-constrained and unconstrained oligo pools

To evaluate sequencing performance of REase-based NinjaSeq (NJ) library preparation, NJ was compared against a standard end-repair/A-tailing workflow adapted for shorter oligo inputs (HP)^9^, followed by ONT sequencing (**Fig. 3a**). Synthetic oligopools were selected to span low complexity (on the order of ≈ 10^3^ distinct oligos) and high complexity (≈ 10^5^ distinct oligos). Moreover, to test whether the presence of the BseGI RRS motif (GGATG) affects sequencing performance, one low-complexity oligopool was generated from payloads encoded with enforced RRS avoidance (w/RRS constraint), whereas the remaining pools were selected from existing oligopools that were designed without RRS constraints (w/o RRS constraint). Design characteristics of oligopools are provided in **Table 1**.

**Figure 3.**
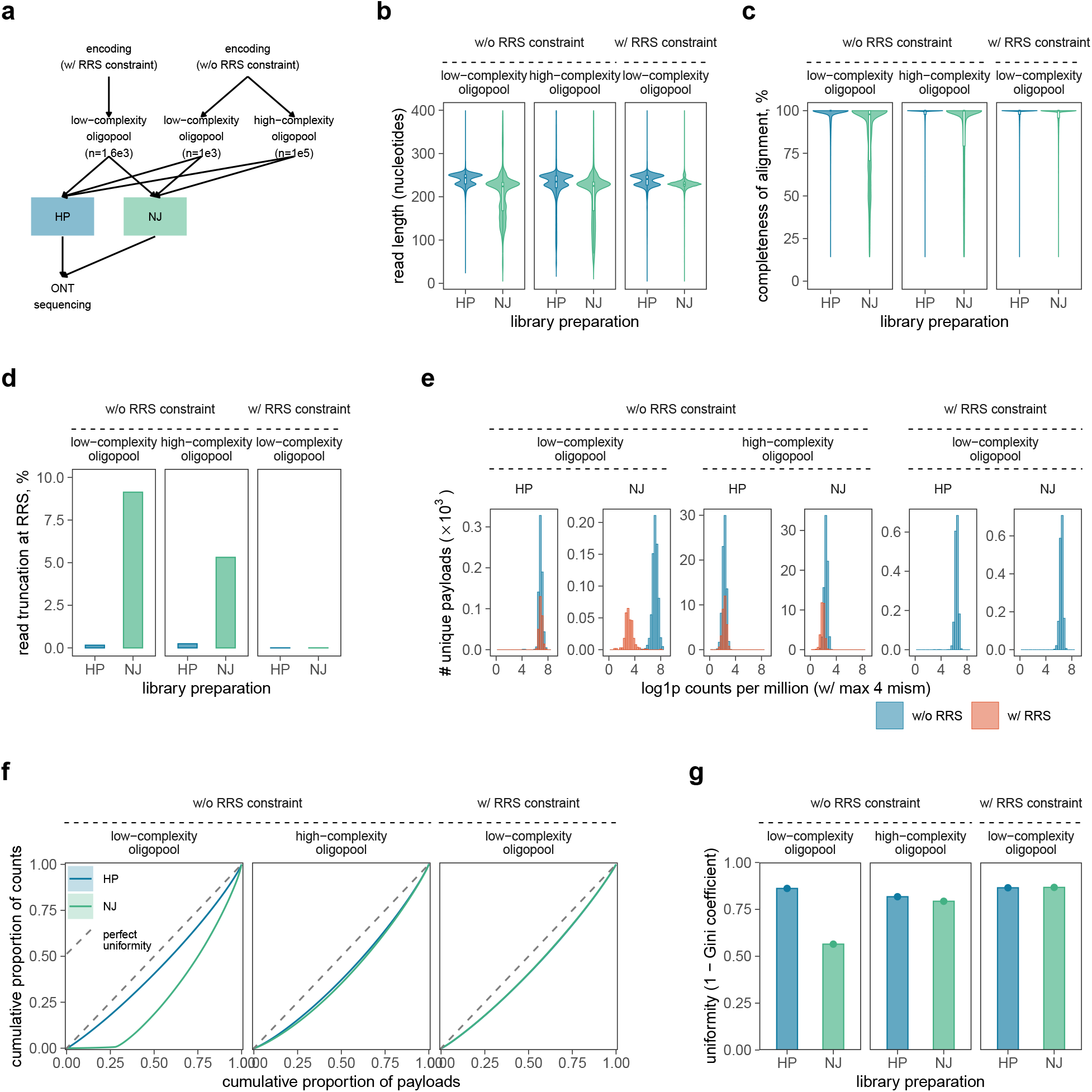
Sequencing performance of REase-based NinjaSeq (NJ) compared with a standard end-repair/dA-tailing workflow (HP) across oligopool complexity and RRS constraint conditions. **a)** Experimental design. Synthetic oligopools were defined by their encoding strategy, with (w/) or without (w/o) RRS constraints for BseGI REase, and by sequence complexity (n). Each pool was processed in parallel using HP or NJ library preparation protocols followed by ONT sequencing. **b)** Read-length distributions for HP and NJ libraries. Violin plots show the density of read lengths, with overlaid boxplots indicating the median and interquartile range. **c)** Completeness of alignment to the expected reference sequences (fraction of read length aligned to reference). **d)** Fraction of reads truncated at the BseGI RRS. **e)** Distribution of per-payload abundance (log1p counts per million; CPM) stratified by whether a payload contains a BseGI RRS (w/RRS vs w/o RRS). **f)** Lorenz curves of payload abundance showing deviations from perfect uniformity (dashed line) for HP and NJ libraries. Shaded ribbons indicate variability based on replicate spread. **g)** Uniformity summary quantified as (1-Gini coefficient) for each condition (higher values indicate more uniform representation across payloads). Each condition represents a single library preparation and sequencing run (N=1).

**Table 1.**
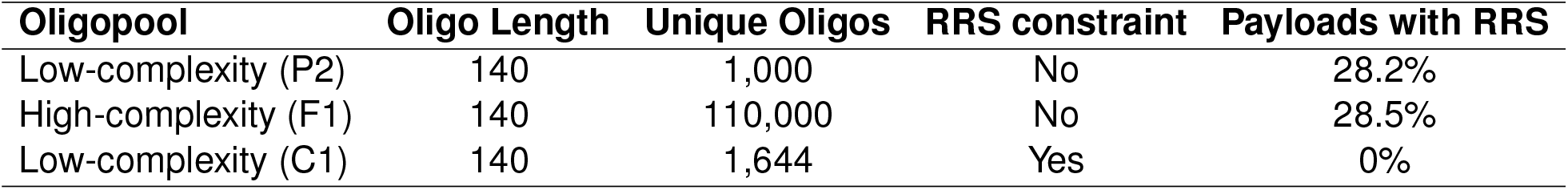
Summary of design parameters and characteristics of the oligopools used in this study. RRS denotes the BseGI restriction recognition site (GGATG).

| Oligopool | Oligo Length | Unique Oligos | RRS constraint | Payloads with RRS |
| --- | --- | --- | --- | --- |
| Low-complexity (P2) | 140 | 1,000 | No | 28.2% |
| High-complexity (F1) | 140 | 110,000 | No | 28.5% |
| Low-complexity (C1) | 140 | 1,644 | Yes | 0% |

Each oligopool was split and processed in parallel using either HP or NJ, followed by ONT sequencing (see **Methods**). This design enabled a head-to-head comparison of library-preparation performance on identical input pools, while also assessing robustness to differences in pool complexity and to the presence or absence of RRS-avoidance constraints.

Examination of read-length distributions revealed marked differences between the workflows (**Fig. 3b**). In oligopools designed without RRS constraints, NJ libraries contained an elevated proportion of reads shorter than the full-length amplicon, suggesting sequence truncation. Conversely, in the low-complexity pool with enforced RRS avoidance, the presence of these truncated species in the NJ library was minimal and comparable to the HP control (**Fig. 3b**). Another observation was that while HP contained two peaks corresponding with single and dual ligated adapters, the single-adapter fraction was notably more pronounced in NJ. The accumulation of singly ligated products in NJ was consistent with observations from the independent mock-adapter ligation experiments (**Fig. 1b**).

The read truncation observed in unconstrained pools directly affected alignment completeness metrics (**Fig. 3c**). While HP libraries consistently yielded high alignment completeness, NJ libraries lacking RRS constraints displayed a broad distribution extending to low completeness values, confirming that the truncated reads spanned only a fraction of the reference sequence (**Fig. 3c**). To identify the source of these partially aligning reads, truncation was quantified specifically at the BseGI recognition motif (**Fig. 3d**). In the absence of RRS constraints, NJ displayed a substantial increase in reads truncated at the RRS compared to HP. This RRS-specific fragmentation was notably more pronounced in the low-complexity oligopool than in the high-complexity oligopool. However, the quality score distributions per-read were similar between NJ and HP libraries (**Supplementary Fig. 2a**), whereas the total sequencing output varied with the composition of the pool and the protocol (**Supplementary Fig. 2b**).

The impact of internal cleavage on library composition was further evident in payload abundance distributions. In the low-complexity pool without RRS constraints, payloads containing the BseGI motif exhibited a downward shift in abundance relative to those lacking the motif (**Fig. 3e**), consistent with the depletion of sequences susceptible to digestion. Consequently, Lorenz curve analysis (**Fig. 3f**) and the derived uniformity metric (1-Gini) (**Fig. 3g**) indicated reduced evenness for NJ in the unconstrained low-complexity pool. In contrast, the high-complexity pool appeared less sensitive to this bias, displaying uniformity metrics that were broadly similar between protocols. Importantly, under the RRS-avoidance constraint, the uniformity of the NJ library improved to match that of HP (**Fig. 3g**). To evaluate potential off-target effects in this constrained context, differential truncation analysis was performed, identifying only rare instances of cleavage (**Supplementary Fig. 2c, d**). These isolated events did not compromise global recovery, demonstrating that RRS-mediated bias can be effectively mitigated through sequence design.

Overall, NJ performance depended on oligopool complexity and RRS content. RRS-associated truncation was most evident in the unconstrained low-complexity pool and was attenuated in the high-complexity pool, despite similar fractions of RRS-containing payloads (**Table 1**). Enforcing RRS avoidance largely eliminated this bias, yielding NJ read completeness and library uniformity similar to HP.

### Decoding performance of NinjaSeq libraries from RRS-constrained and unconstrained oligo pools

To assess the impact of NinjaSeq (NJ) library preparation on data retrieval, decoding performance was evaluated by analyzing error rates and reconstruction accuracy across the RRS-constrained and unconstrained oligopools. All tested pools were synthesized using a design structure similar to the DNAformer encoder^10^, comprising a 12-base index for unique identification and a 128-base payload containing the encoded information and error-correction redundancy. The resulting 140-nt constructs were flanked by primers for amplification, which were trimmed from the sequencing reads prior to analysis. While the two low-complexity pools (w/o and w/RRS-constraint) encoded random information, the high-complexity pool (w/o RRS-constraint) encoded both random and semantic files^10^.

First, sequencing error characteristics were quantified using the SOLQC tool^11^, which bins reads by index and aligns them to the reference design to estimate base-level errors (deletions, substitutions, and insertions). Analysis of error distributions revealed that NJ consistently exhibited higher error rates than the standard high-performance (HP) workflow at both the per-read and per-cluster levels (**Fig. 4a,b**). Specifically, in the RRS-unconstrained low-complexity pool, the mean error-rate difference between NJ and HP was marked, reaching 0.027 per read (0.118 vs. 0.091) and 0.078 per cluster (0.114 vs. 0.036). In the high-complexity pool, these differences narrowed to 0.012 per read and 0.012 per cluster. Importantly, the RRS-constrained pool showed intermediate differences of 0.016 per read and 0.011 per cluster. This trend indicates that the additional error burden introduced by NJ depends on pool complexity and motif content. When clusters containing RRS motifs were computationally filtered from the unconstrained datasets, the error-rate gap between NJ and HP decreased substantially (to approx. 0.001–0.01 at the cluster level), confirming that enforcing RRS avoidance can significantly improve the base-level accuracy of NJ-based sequencing.

**Figure 4.**
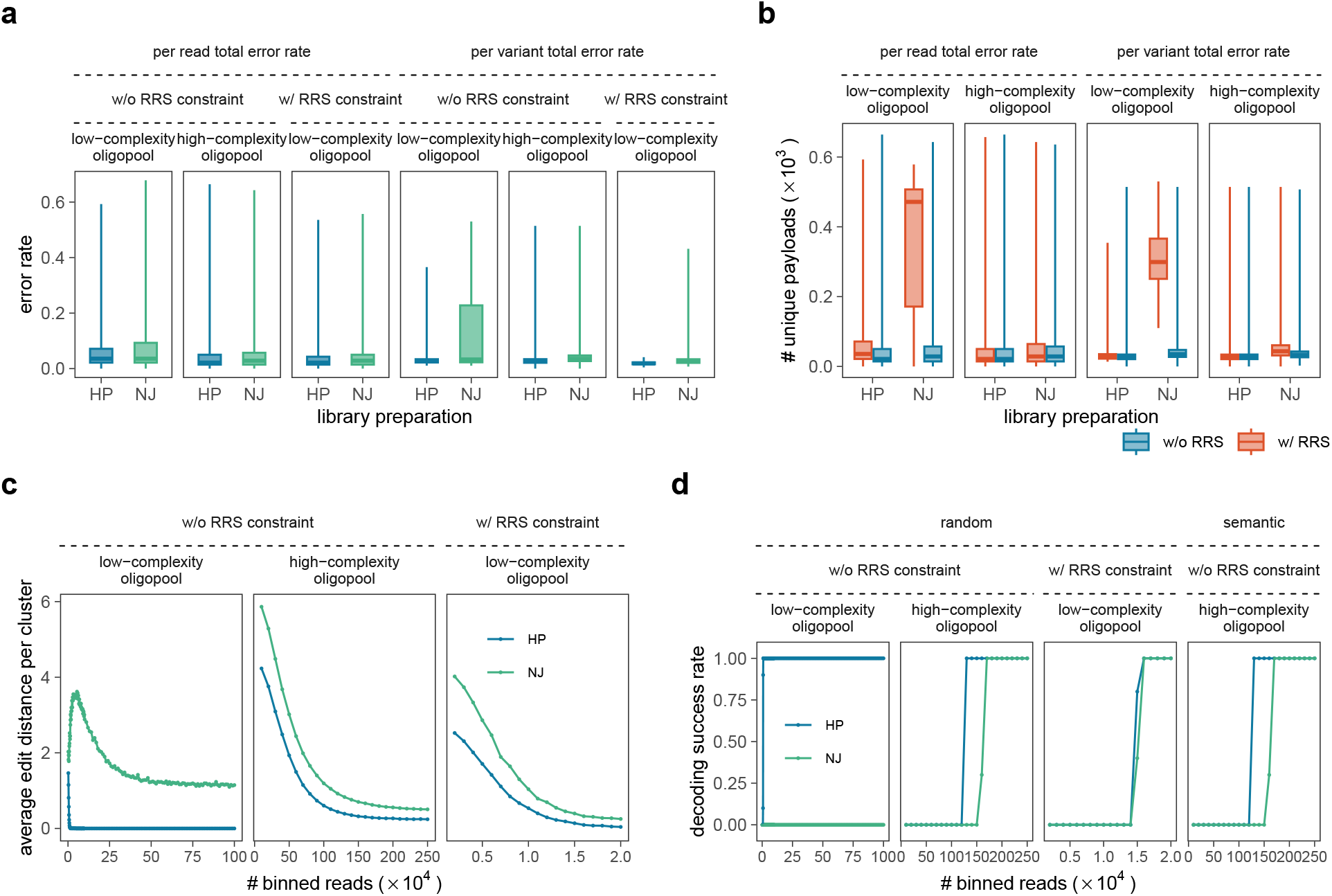
Evaluation of Decoding performance of NinjaSeq Libraries. **a)** Error rate analysis. The figure shows a boxplot with the error rates distribution across the sequenced libraries, stratified by the library prep method. The error rates are shown at the read level, and at the cluster level (per variant). **b)** Error rate analysis stratified by RRS constraints. The boxplots show the distribution of error rates across sequenced libraries, partitioned by the presence or absence of RRS constraints. Results are shown for both read-level error rates and cluster-level (per-variant) error rates, and are further stratified by the library preparation method. **c)** Evaluation of the edit distance of the reconstructed oligos as a function of the number of binned reads. The values are shown per oligo pool and per library preparation method. **d)** The decoding success rate as a function of the number of binned read. Each condition represents a single library preparation and sequencing run (N=1).

To determine how these error profiles affect information recovery, a decoding pipeline was applied using the CPL reconstruction algorithm^10,12^. Because reconstruction success depends on both the error rate and the coverage depth, datasets were downsampled to varying numbers of binned reads (**Fig. 4c,d**). Comparison of the reconstructed sequences against the original designs showed that the average edit distance decreased as the number of binned reads increased (**Fig. 4c**). Although NJ libraries initially yielded larger edit distances than HP, this performance gap narrowed at higher coverage depths.

Finally, decoding success rates were measured over 20 independent downsampling experiments (**Fig. 4d**). Across all conditions, a minimum threshold of binned reads was required to achieve successful decoding. This threshold was higher for NJ than for HP, reflecting the increased error rates associated with the REase-based workflow. However, for the RRS-constrained pool, the coverage thresholds necessary for successful decoding were virtually identical between NJ and HP, in distinct contrast to the RRS-unconstrained low-complexity pool, where decoding proved more challenging. These observations indicate that while NJ libraries exhibit higher per-base error rates than standard end-repair/dA-tailing, the impact on decoding can be mitigated by RRS-constrained sequence design and adequate sequencing coverage.

### Random access capabilities of NinjaSeq during library preparation

To demonstrate the random access capability of NinjaSeq, a selective retrieval experiment was performed using a mixture of three distinct oligopools (Files A, B, and C) combined at defined relative abundances proportional to their unique payload counts (File A: 0.37; File B: 0.23; File C: 0.40). In this design, only the target file (File A) contained the specific recognition site (RRS) at its ends. While Files B and C were susceptible to random internal cleavage, they lacked the terminal sites necessary to generate valid library constructs. Consequently, REase digestion was expected to yield full-length, adapter-compatible products exclusively for File A (**Fig. 5a**). To benchmark this selectivity, the same mixture was processed in parallel using standard end-repair/dA-tailing (HP), which prepares all DNA ends for sequencing indiscriminately.

**Figure 5.**
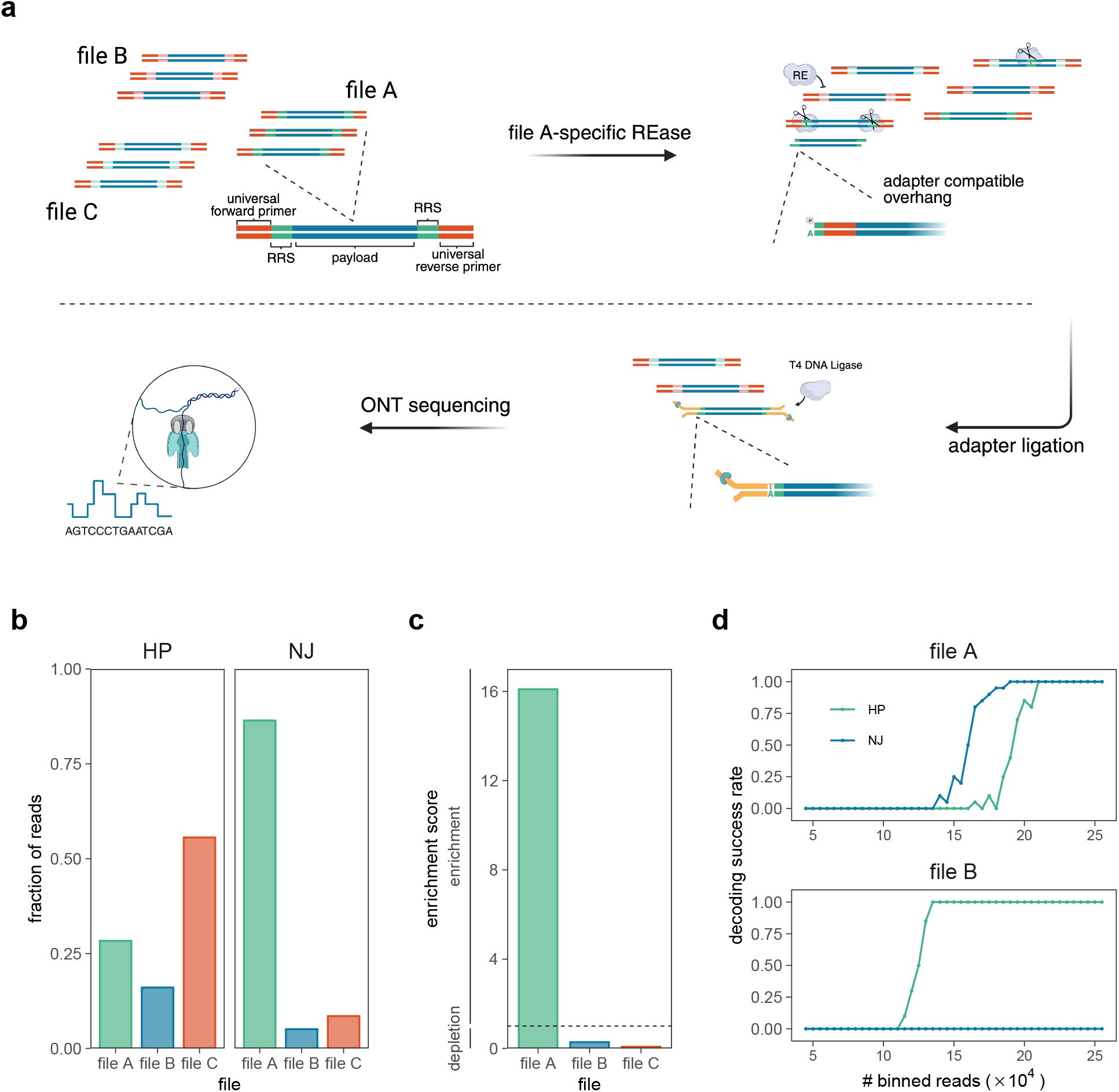
Evaluation of NinjaSeq random access capabilities. **a)** Schematic of the selective retrieval strategy. Unique restriction recognition sites (RRS) serve as file-specific addresses. In a mixture of three files (A, B, and C), the addition of a File A-specific REase targets the flanking RRSs, generating ligation-compatible termini exclusively at the ends of the target. T4 DNA ligase subsequently attaches sequencing adapters to these specific overhangs, enabling the recovery of the full-length File A oligo, while non-target files (B and C) do not yield functional library constructs. **b)** Fraction of mapped reads assigned to the target (File A) versus background files (B and C). Libraries prepared using the standard end-repair/dA-tailing protocol (HP) exhibit a distribution consistent with the input mixture (File A: 0.37; File B: 0.23; File C: 0.40), whereas the NinjaSeq (NJ) library displays a distinct shift in specificity, enriching the target File A to 86.4% of total reads. **c)** Enrichment scores calculated as the odds ratio of observing a specific file in the NJ library relative to the HP control. The dashed line indicates a neutral ratio (1). The target File A exhibits a 16.1-fold enrichment, while non-target files B and C are effectively depleted (odds ratios of 0.28 and 0.07, respectively). **d)** Decoding success rate as a function of sequencing depth (number of binned reads) for the retrieved target (File A) and the background (File B). The target file achieves 100% recovery (decoding rate = 1.0) in the NinjaSeq library with comparable efficiency to the control, while the depleted background files do not reach sufficient coverage for decoding. Each condition represents a single library preparation and sequencing run (N=1). Created with BioRender.com.

Following sequencing, reads were adapter-clipped and size-selected (minimum length 180 nt; see Methods) to remove truncated fragments and off-target cleavage products. As expected, the standard workflow (HP) captured all three files, with their relative abundances reflecting the composition of the input mixture. In contrast, NinjaSeq (NJ) substantially altered this distribution, enriching the target File A to 86.4% of the total reads (**Fig. 5b**). Relative to the non-selective control, this corresponds to a 16.1-fold enrichment (odds ratio) of the target file in this proof-of-concept experiment (**Fig. 5c**). Conversely, Files B and C were effectively depleted (odds ratios: 0.28 and 0.07, respectively). To verify data integrity, we analyzed the decoding performance of the retrieved sequences (**Fig. 5d**). The target File A achieved 100% decoding success in the NinjaSeq condition, reaching saturation at approximately 190,000 reads. In contrast, the non-targeted File B failed to decode under the same conditions, demonstrating that the enzymatic selection effectively filters out background data. These data confirm that NinjaSeq enables specific physical retrieval of a target file from a heterogeneous background without the need for additional purification steps.

## DISCUSSION

This work presents NinjaSeq, a streamlined library preparation workflow for ONT nanopore sequencing that leverages restriction endonucleases (REases) to bypass standard end repair and dA tailing. The results demonstrate that by effectively constraining the sequence design to avoid recognition sites, REase based preparation achieves decoding performance comparable to standard end repair and dA tailing protocols.

The utilization of REases in this context represents a conceptual revival of “legacy” molecular tools in the era of programmable gene editing. While the last decade has been defined by the rise of programmable nucleases such as CRISPR-Cas9, which can be targeted to arbitrary sequences^13^, REases have traditionally been viewed as rigid tools limited by fixed recognition motifs. However, the paradigm of DNA data storage fundamentally inverts this constraint. In this domain, the data carrier itself is synthetic and fully programmable. Consequently, rather than requiring an enzyme that adapts to a fixed biological target, the target sequences can be adapted to match the specific requirements of highly characterized, robust, and commercially available enzymes. The results confirm that when the programmability is shifted from the enzyme to the information encoding, specifically through the use of RRS constrained encoding, the rigidity of REases becomes an asset, offering high specificity and predictability.

The primary advantage of this approach is the democratization and simplification of the sequencing workflow. The vast diversity of cataloged REases^14^ provides a rich toolkit of enzymes with varying overhang characteristics and thermal profiles, offering flexibility that standard end repair mixes lack. Furthermore, REases are widely available, stable, and less expensive than the proprietary enzyme mixes required for dA tailing. A reagent-by-reagent cost comparison of the end-preparation step is provided in **Supplementary Table** 3. Based on current list prices, NinjaSeq end-preparation reagents are approximately 30-fold less expensive per reaction than the corresponding commercial end-repair/dA-tailing module. This cost reduction is important for the scalability of DNA data storage, where the cost of reading data remains a significant bottleneck. Moreover, the feasibility of a one-pot reaction where digestion and ligation occur simultaneously was demonstrated. While the current data indicates that the two step protocol remains more efficient, the compatibility of these enzymatic steps paves the way for fully automated, liquid handling friendly workflows that minimize user intervention and reagent waste.

The performance of NinjaSeq was shown to be intrinsically linked to both the presence of RRS motifs within the payload and the overall diversity of the oligonucleotide pool. Analysis revealed that unconstrained pools suffer from significant read truncation and coverage bias, particularly in the low-complexity oligo pool, where RRS specific depletion leads to elevated error rates and decoding failure. However, by modeling RRS avoidance as a constrained coding problem on a graph, it was shown that these off target effects can be virtually eliminated. Importantly, the theoretical capacity loss incurred by these constraints is negligible for typical recognition site lengths (4 to 6 bp), allowing for robust library preparation without significantly penalizing information density. When these constraints were applied, the decoding performance of NinjaSeq, measured by error profiles and required coverage, converged with that of the standard end repair and dA tailing workflow.

Beyond the library preparation, the specificity of REases opens new avenues for functional data retrieval, such as random access. Current random-access methods typically rely on PCR amplification using specific primers to retrieve a file of interest from a pool^15^. NinjaSeq demonstrates an alternative or complementary strategy: by encoding different files or data blocks with distinct flanking RRS motifs, specific datasets can be chemically selected for sequencing library preparation without amplification bias. By using a specific REase to activate only the target subset of the pool for adapter ligation, physical random access is achieved during the library preparation phase itself. A similar, albeit multi-step, approach was recently described using a combination of Cas9 and dA tailing enzymes^16^.

In conclusion, NinjaSeq illustrates the practical benefits of integrating digital sequence design with physical library preparation. The findings demonstrate that by incorporating defined constraints into the encoding scheme, the necessity for standard end repair and dA tailing is removed, yielding a workflow that is both cost efficient and robust. This approach not only reduces the cost and complexity of DNA data storage workflows but also highlights the potential of revisiting foundational molecular biology tools to solve modern challenges in synthetic biology.

### Limitations of the study

Several limitations of NinjaSeq warrant consideration. First, the current workflow relies on REases that generate 1–3 nt 3′-A overhangs, limiting the range of compatible enzymes relative to fully programmable end-preparation chemistries. Second, although the RRS-constrained encoder preserves nearly the full encoding capacity at typical recognition-site lengths, motif selection is not completely orthogonal across the sequence catalogue, requiring coordinated design of payload sequences and primer-flanking regions. Third, NJ libraries exhibited modestly higher per-base error rates than HP libraries within the RRS-constrained pool. The basis of this effect remains unclear but may reflect altered motor-protein behaviour in the vicinity of the BseGI scar. Fourth, the current one-pot restriction–ligation workflow is less efficient than the corresponding two-pot implementation, indicating scope for further optimisation. Finally, the present study represents a proof of concept demonstrated in a single experimental dataset. Additional protocol-level replication, together with orthogonal random-access experiments employing multiple REases to selectively retrieve different files from the same pool, will be required to establish the robustness, scalability, and general applicability of the approach.

## METHODS

### Scope and replication

This study presents NinjaSeq as a methodological demonstration. Unless otherwise stated, each library preparation and sequencing condition was performed once (N=1). Variability within each experiment was assessed across per-payload distributions (thousands of unique oligos per pool), rather than across protocol-level replicates. Statistical comparisons between library-preparation protocols were therefore not performed, and all between-protocol differences are reported as descriptive observations.

### Exploratory assessment of REase-based end-preparation and ligation

#### REase selection for end-preparation

For the exploratory experiments, REases were selected according to several key criteria. First, the enzymes had to produce 3′ overhangs after cleavage. Second, the RRS needed to contain a programmable 3′ region (having Ns), allowing the overhang to include at least one (-A) motif. Additional selection criteria included fast digestion kinetics, distinct recognition patterns (non-palindromic), and the ability to generate varying numbers of overhangs per cleavage event. Based on these criteria, a subset of REases was selected for experimental validation: MnlI, BseGI, AasI, AdeI, and BstXI (see **Table 2** for characteristics).

**Table 2.**
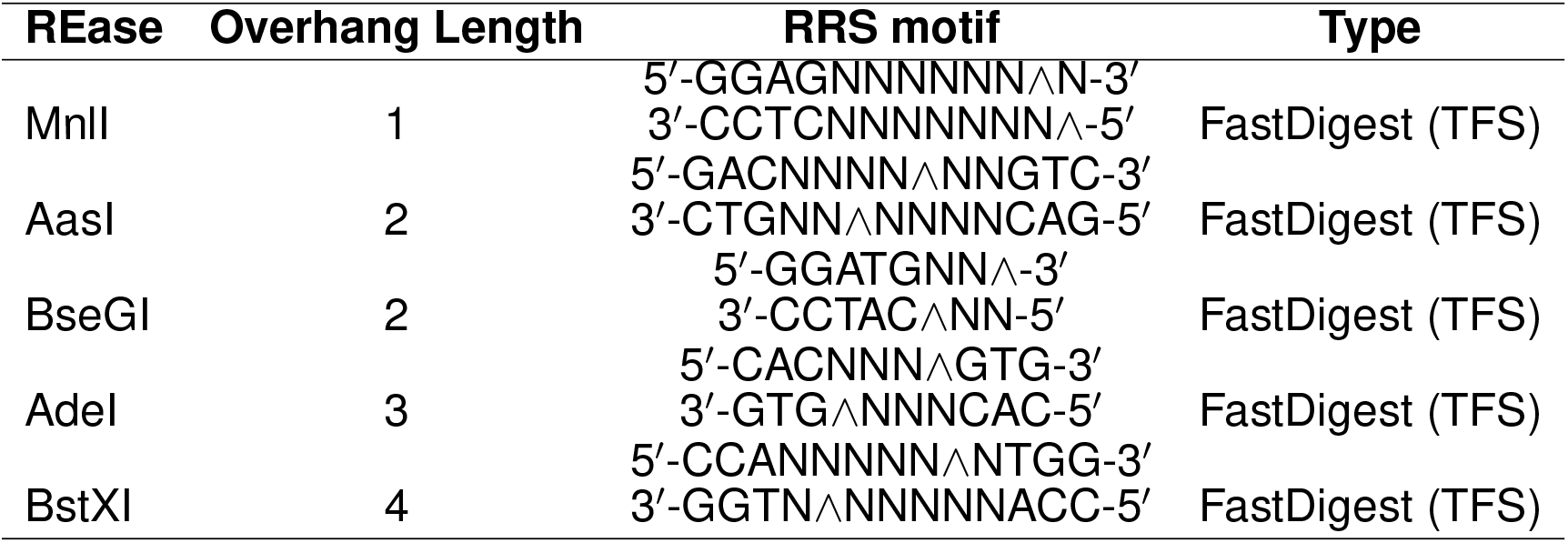
Characterization of REases used in the exploratory assessment of end-preparation for ligation.

| REase | Overhang Length | RRS motif | Type |
| --- | --- | --- | --- |
| MnlI | 1 | 5'-GGAGNNNNNNN^N-3'<br>3'-CCTCNNNNNNN^5' | FastDigest (TFS) |
| AasI | 2 | 5'-GACNNNN^NNGTC-3'<br>3'-CTGNN^NNNNCAG-5' | FastDigest (TFS) |
| BseGI | 2 | 5'-GGATGNN^3'<br>3'-CCTAC^NN-5' | FastDigest (TFS) |
| Adel | 3 | 5'-CACNNN^GTG-3'<br>3'-GTG^NNNCAC-5' | FastDigest (TFS) |
| BstXI | 4 | 5'-CCANNNNN^NTGG-3'<br>3'-GGTN^NNNNNACC-5' | FastDigest (TFS) |

#### Oligo and primer design

To introduce RRS into the experimental constructs, a 120-nt template sequence (5′-TGCATCAT CTCTACACATAGCTGCGAGTACGCATCTACAGATCTCGTGTGTATGACAGATCGACGATG ATGCGAGATCACGCGTACAGCATCATGACTGCGTGTATCGTCGTAGCGAGTC-3′; Eurofins) was amplified by PCR using specifically designed primers. Each primer was designed according to uniform logic. The primer design incorporated three key elements: (1) additional nucleotides (NNN) at the 5′ end to facilitate proper enzyme binding; (2) the specific RRS for the target enzyme; and (3) a 3′ terminal overhang of varying length (1-4 nucleotides) terminating in -T (or -C in specific cases). Following RE digestion, the resulting overhangs consisted of 1 to 4 adenine (-A) nucleotides. Complete primer sequences are provided in **Supplementary Table 1**.

#### PCR amplification

To introduce RRS into the template DNA, PCR was performed using Kapa HiFi DNA polymerase (Roche) according to the manufacturer’s guidelines. Each 50 *µ*l reaction contained: 25 *µ*l Kapa HiFi master mix, 1.5 *µ*l forward primer (10 *µ*M), 1.5 *µ*l reverse primer (10 *µ*M), 1 ng template DNA, and nuclease-free water to volume. Thermal cycling conditions consisted of an initial denaturation at 95°C for 3 min, followed by 17 cycles of 98°C for 20 s, primer-specific melting temperature (Tm; **Supplementary Table 1**) for 10 s, and 72°C for 10 s, with a final extension at 72°C for 1 min.

Following amplification, PCR products were purified using the Monarch PCR & DNA Cleanup Kit (5 *µ*g capacity, NEB) according to the manufacturer’s protocol. The DNA to DNA binding buffer ratio was maintained at 1:5 (v/v). Purified DNA concentrations were quantified using the Qubit dsDNA HS Assay Kit (TFS).

#### Restriction digestion

To generate the desired overhangs, purified PCR products (150 *×* ng) were digested with the appropriate restriction endonuclease. Each 20 *µ*l reaction contained: 2 *µ*l Fast Digest buffer (10×, TFS), 1 *µ*l restriction enzyme, DNA template (150 ng), and nuclease-free water to volume. Enzyme-specific incubation and thermal inactivation conditions are provided in Supplementary 2. Following digestion, reactions were purified using the Monarch PCR & DNA Cleanup Kit (5 *µ*g capacity, NEB) with a 1:5 (v/v) DNA to binding buffer ratio. Purified DNA concentrations were quantified using the Qubit dsDNA HS Assay Kit (TFS).

#### Mock adapter ligation

To mimic double-stranded DNA (dsDNA) part of ONT sequencing adapters, synthetic dsDNA adapters were generated by annealing two complementary oligonucleotides matching dsDNA ONT adapter sequences. Oligonucleotides were synthesized using a Kilobaser DNA printer (Kilobaser): top strand (5′-CCTGTACTTCGTTCAGTTACGTATTGCT-3′) and bottom strand (5′-GCAATACGTAACTGAACGAAGTACAGG-3*′*).

Adapter annealing reactions (10 *µ*l total volume) contained: 1 *µ*l top strand oligonucleotide (100 *µ*M), 1 *µ*l bottom strand oligonucleotide (100 *µ*M), 1 *µ*l T4 DNA ligase buffer (10 *×*), and 7 *µ*l nuclease-free water. Annealing was performed by heating to 95°C for 5 min, followed by gradual cooling at 5°C per minute (90°C for 1 min, 85°C for 1 min, etc.) until reaching 25°C. Annealed adapter concentrations were quantified using the Qubit dsDNA HS Assay Kit (TFS).

As a positive control, the 120-nt template oligonucleotide underwent standard end-preparation, including adenylation and phosphorylation, using the NEBNext Ultra II End Repair/dA-Tailing Module (NEB). The reaction (60 *µ*l) contained: 200 fmol DNA template, 7 *µ*l End-Prep Reaction Buffer (10 *×*), 3 *µ*l End-Prep Enzyme Mix, and nuclease-free water to volume. The reaction was incubated at 20°C for 5 min, followed by 65°C for 5 min.

Adapter ligation was performed in 100 *µ*l reactions containing: 30 ng DNA substrate, 25 *µ*l T4 DNA ligase buffer (10×), 10 *µ*l T4 DNA ligase, adapters at a 7:1 molar ratio (adapter: DNA; e.g., 49 ng adapters for 30 ng DNA), and nuclease-free water to volume. Reactions were incubated at room temperature for 20 min. Following ligation, samples were purified using SPRI magnetic beads (Beckman Coulter) at a 1.8:1 bead-to-DNA ratio with isopropanol to retain adapter-ligated products. Briefly, beads were mixed with 270 *µ*l isopropanol prior to sample addition, followed by standard bead purification protocols. Purified DNA concentrations were quantified using the Qubit dsDNA HS Assay Kit (TFS). Ligation product size distributions were assessed on an TapeStation 2200 (Agilent) with D1000 ScreenTape.

#### Forbidden-pattern constrained codes

To mitigate internal cleavage of payload-encoding DNA strands by REases, designed sequences can be constrained to avoid specific forbidden patterns (such as REase recognition sites). This approach naturally necessitates the use of constrained codes. The definitions of constrained systems, their graph presentations, and the characterization of capacity via the spectral radius are standard and follow the treatment in^17^.

#### Definitions

Let Σ denote a finite alphabet, and let *ℱ* ⊂ Σ^+^ be a finite set of substrings that are not allowed to appear in the designed sequence.

##### Definition 1.

*The* forbidden-pattern constrained code *associated with F is defined as*

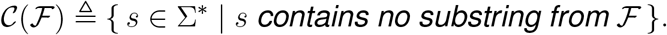

Using constrained codes reduces the probability that error-inducing patterns appear in the sequence, thereby decreasing the error rate of our protocol. However, enforcing constraints necessarily limits the information that can be encoded. This tradeoff is quantified through the capacity of the constrained system.

##### Definition 2.

*The* capacity *of the constrained system 𝒞* (*ℱ*) *is defined as*

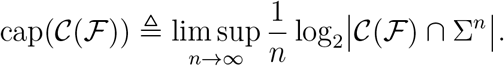

#### Graph construction

To compute the capacity and later integrate the constraint into our reconstruction pipeline, we build a finite labelled directed graph that generates exactly all sequences avoiding the patterns in *ℱ*. We let *m* = max_*f*∈*F*_ |*f* | denote the length of the longest forbidden pattern.

##### Definition 3.

*The* constraint graph *associated with, denoted G*_*F*_ = (*V, E, L*), *is constructed as follows:*

- *Each vertex v* ∈ *V corresponds to a distinct prefix of length at most m* − 1 *that does not contain any substring from F*.
- *For any u, v* ∈ *V and a* ∈ Σ, *we include a directed edge u → v labelled by a if appending a to the string represented by u produces the string represented by v, and this extension does not introduce any forbidden pattern*.
- *The edge-labelling function L* : *E →* Σ *assigns to each edge the symbol a used in the transition*.

By construction, every directed path in *𝒢*_*ℱ*_ corresponds to a unique sequence in *𝒞* (*ℱ*), and every sequence in *𝒞* (*ℱ*) corresponds to at least one path in the graph. Thus, *G*_*F*_ provides an ordinary and lossless presentation of the constrained system.

Let *A*_*ℱ*_ denote the adjacency matrix of *𝒢*_*ℱ*_, where *A*_*ℱ*_ (*i, j*) counts the number of edges from vertex *i* to vertex *j*. For such a graph, the capacity of the constrained system is given by

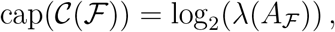

where *λ*(*A*_*ℱ*_) is the spectral radius of *A*_*ℱ*_.

In our application, certain substrings cause systematic errors not only when they appear in the designed sequence, but also when their reverse-complement appears. To enforce this additional constraint, for every forbidden pattern *f* ∈ *ℱ* we also consider its reverse-complement RC(*f*). Since the reverse-complement of a pattern may induce the same type of errors in the NinjaSeq protocol, we simply append RC(*f*) to the forbidden set. Thus, the augmented set of forbidden patterns is

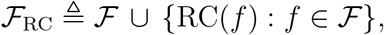

and all definitions above apply unchanged when *ℱ* is replaced by *ℱ*_RC_.

#### Algorithmic construction of the constraint graph

Although the definition of the constraint graph *𝒢*_*ℱ*_ is conceptual, it is useful to describe the explicit procedure we employ to construct it from a given alphabet Σ and forbidden set *ℱ* (including reverse-complements when applicable). The algorithm operates at a high level in three stages: (1) identifying all valid prefix states, (2) determining valid state transitions, and (3) assembling the adjacency matrix.

##### State generation

We first generate the set of graph vertices by collecting all distinct prefixes of all forbidden patterns, including the empty prefix. These prefixes represent all possible *states* the system may be in before emitting the next symbol, and ensure that the graph captures all contexts relevant to detecting forbidden patterns.

##### Transition determination

For each state *s* and for each letter *a* ∈ Σ, we determine whether the extension *sa* introduces a forbidden substring. If the extension is valid, we compute the next state by identifying the longest suffix of *sa* that is itself a valid prefix state. This ensures that the graph transition preserves all contextual information needed to avoid forbidden patterns.

##### Adjacency matrix construction

Once all transitions are determined, we construct the adjacency matrix *A*_*F*_, where each entry counts the number of allowed edges between states. This matrix provides a complete representation of the constrained system, enabling capacity computation through its spectral radius.

###### Algorithm 1

High-Level Construction of the Constraint Graph

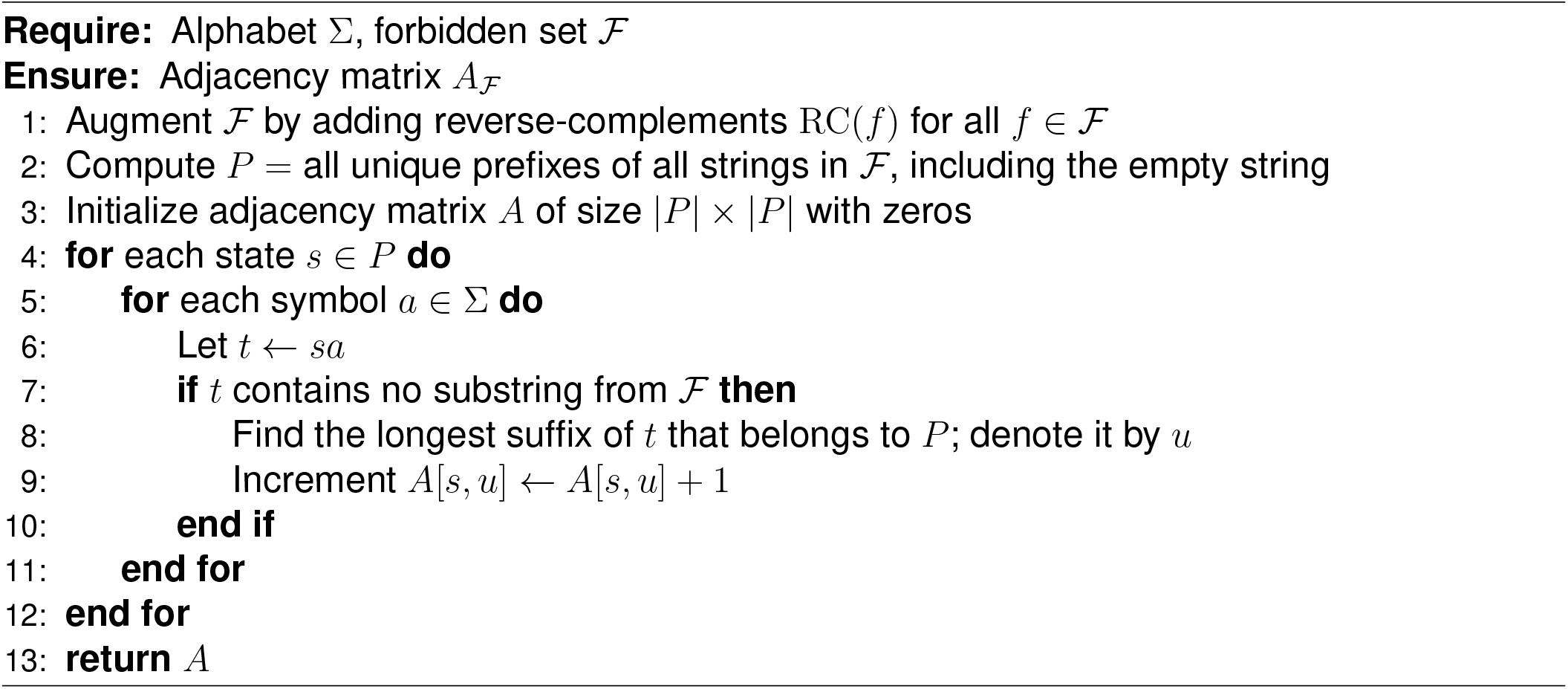

### Sequencing and decoding performance of NinjaSeq protocol

#### Oligo pool design

The oligos used in all pools were designed and encoded using the DNAformer encoder^10^. Each oligo consists of a 12-base index, a 128-base payload (which includes both the information and the redundancy used for error correction and mitigation), and flanking primers (34 and 33 bases), resulting in a total length of 207 bases. In general, the overall information density of the scheme is 1.6 bits per base.

The 128 bases of payload were encoded using an encoder which includes error-correcting codes, based on tensor-product^18^ code and Reed Solomon^19^ code, and constrained coding techniques. The error-correction capability of the code ensures that the stored data can be retrieved even if up to 6.25% of the oligos have some errors. The structural constraint of the code enforces each of the sequences to have almost balanced GC content, specifically between 42.5% and 57.5%. Additionally, the maximum homopolymer length is 6. For the RRS constrained pool, oligos were generated by a constrained encoder that excludes the RRS motif (and its reverse complement) from the payload. More details about the error correction technique, together with an implementation of the encoder and decoder used in this study can be found in^10^.

#### NinjaSeq library preparation and ONT sequencing

##### PCR amplification (NinjaSeq)

For NinjaSeq library preparation, the oligo pools were amplified separately using universal primers containing BseGI restriction recognition sites (RRS) at the 5′ end. The BseGI recognition site generates a programmable 3′ overhang upon digestion (5′-GGATGNN↓-3′/3′-CCTAC↑NN-5′). Primers were designed as follows: forward primer, 5′-TC AGGATGTTTCGTCGGCAGCGTCAGATGTGTATAAGAGACAG-3′; reverse primer, 5′-TCAGGATGTTGTCTCGTGGGCTCGGAGATGTGTATAAGAGACAG-3′. PCR amplification was performed using Kapa HiFi DNA polymerase (Roche) according to the manufacturer’s guidelines. Each 50 *µ*l reaction contained: 25 *µ*l Kapa HiFi master mix, 0.75 *µ*l forward primer (10 *µ*M), 0.75 *µ*l reverse primer (10 *µ*M), 1 ng template DNA (pre-denatured at 98°C for 5 min), and nuclease-free water to volume. Thermal cycling conditions consisted of an initial denaturation at 95°C for 3 min, followed by 17 cycles of 98°C for 20 s and 72°C for 10 s with a final extension at 72°C for 1 min. Following amplification, PCR products were purified using the Monarch PCR & DNA Cleanup Kit (5 *µ*g capacity, NEB) with a 1:5 (v/v) DNA to binding buffer ratio, and concentrations were quantified using the Qubit dsDNA HS Assay Kit (TFS).

##### BseGI digestion

Purified amplicons (150 ng) were then digested with BseGI restriction endonuclease. Each 20 *µ*l digestion reaction contained: 2 *µ*l Fast Digest buffer (10*×*, TFS), 1 *µ*l BseGI, DNA template (150 ng), and nuclease-free water to volume. The reaction was incubated at 37°C for 5 min, followed by thermal inactivation at 80°C for 5 min. Post-digestion samples were purified using the Monarch PCR & DNA Cleanup Kit with a 1:5 (v/v) DNA to binding buffer ratio, and concentrations were determined using the Qubit dsDNA HS Assay Kit.

##### Adapter ligation

Purified, digested DNA (250 fmol) was then subjected to adapter ligation following the ONT standard protocol using the SQK-LSK114 ligation sequencing kit. Ligation reactions contained: 250 fmol digested DNA, 25 *µ*l LNB buffer, 10 *µ*l T4 DNA ligase, and 5 *µ*l sequencing adapters. After thorough mixing, reactions were incubated at room temperature for 20 min. Following ligation, samples were purified using SPRI magnetic beads (Beckman Coulter) at a 1.8:1 bead-to-DNA ratio following standard protocols, and final concentrations were quantified using the Qubit dsDNA HS Assay Kit. A detailed step-by-step NinjaSeq protocol is available at protocols.io (https://dx.doi.org/10.17504/protocols.io.5jyl8xr9rv2w/v1).

##### HP control library preparation

As a control, the same oligonucleotide pools underwent standard end-repair/A-tailing library preparation using the High-Performance protocol^9^ with the SQK-LSK114 kit. Control pools were amplified using standard universal primers without RRS: forward primer, 5′-TCGTCGGCAGCGTCAGATGTGTATAAGAGACAG-3′; reverse primer, 5′-GTCTCGT GGGCTCGGAGATGTGTATAAGAGACAG-3′. PCR amplification was performed as described above, with the following modifications: each 50 *µ*l reaction contained 1 *µ*l of each primer (10 *µ*M) instead of 0.75 *µ*l. Thermal cycling conditions and purification procedures were identical to those used for NinjaSeq samples. Purified control samples (250 fmol dsDNA) were then subjected to end-preparation and adapter ligation following the High-Performance protocol. Briefly, DNA was phosphorylated at the 5′ end and adenylated at the 3′ end according to the manufacturer’s instructions. Without intermediate purification, ligation adapters were attached using the Quick T4 DNA Ligation Module (NEB) with an extended incubation time of 20 min at room temperature. Following ligation, samples were purified using AMPure XP beads (Beckman Coulter) at a 1.8:1 bead-to-DNA ratio and eluted. The final DNA concentrations were measured using the Qubit dsDNA HS Assay Kit (TFS).

##### ONT sequencing

All libraries (both NinjaSeq and control) were sequenced using ONT Prome-thION or MinION flow cells. For each sequencing run, approximately 90 and 100 fmol of the prepared library were loaded, respectively. The oligo pools were sequenced for 24 hours, followed by downstream sequencing analysis.

##### Sequencing data processing and bioinformatics analysis

Raw ONT signal data collected in POD5 format were basecalled using Dorado v1.1.1 with the simplex SUP model (dna r10.4.1 e8.2 400bps sup@v5.0.0). Basecalled reads were converted from unaligned BAM (uBAM) to FASTQ format using SAMtools^20^ (v1.13). For random access experiment libraries, reads were additionally preprocessed using cutadapt^21^ (v5.2) to remove adapters and filter by length. The 5′ adapter sequence was trimmed with the following parameters: `-g CCTGTACTTCGTTCAGTTACGTATTGCT --overlap 8 --error-rate 0.25 --revcomp--discard-untrimmed --minimum-length 180`. Reads were then aligned to reference payload sequences using minimap2^22^ (v2.30) with the following parameters: `-ax sr -L --MD -t8 -Y--eqx -k10 -w5 -m10 --secondary=yes`. To quantify reference payload sequences, aligned reads were filtered based on quality and alignment metrics: only reads with quality scores ≥ 9, ≤ 4 mismatches (NM ≤ 4), and aligned lengths of 140 ± 4 bp were retained and counted. Pay-load sequence uniformity was evaluated using Lorenz curves and the Gini coefficient, which was calculated from the cumulative distribution of mapped read counts across all reference sequences. The Gini coefficient ranges from 0 (perfect uniformity, where all sequences have equal coverage) to 1 (maximum inequality, where coverage is concentrated in a single sequence), and was computed as *G* = 1 *−* 2*A*, where *A* is the area under the Lorenz curve. For intuitive interpretation, uniformity scores were reported as *U* = 1 *− G*, where higher values indicate more uniform coverage. Data analysis and visualization were performed using custom Python (v3.12.9) and R (v4.5.1) scripts, which are available in the GitHub repository at https://github.com/genomika-lt/NinjaSeq.

##### Error Characterization

Error behavior analysis was performed using the SOLQC tool^11^. The tool uses the set of predefined indices in order to bin the sequencing reads based on their original encoded sequences. Following that, alignment is performed on each read to align it with its corresponding designed encoded sequence to assess the number of insertion, deletion, and substitution errors. The tool reports error rates at the cluster level and at the read level.

##### Decoding

The decoding pipeline consists of three main stages: binning, reconstruction, and error-correcting decoding.

In the binning stage, reads are grouped using a predefined set of indices. Specifically, the index region of each read is inspected, and reads that share the same index are assigned to the same bin, corresponding to a single designed sequence. Due to the presence of sequencing errors in the index region, some reads may fail to match any valid index and are therefore filtered out during this step.

Following binning, sequence reconstruction is performed using the CPL reconstruction algorithm^10^. This algorithm applies a dynamic programming approach to estimate the original encoded sequence from a cluster of noisy reads. The computational complexity of the CPL reconstruction is quadratic in the sequence length.

Finally, the DNAformer error-correcting decoder is applied to the reconstructed sequences to correct any remaining errors. To evaluate the effect of sequencing coverage on decoding performance, we performed downsampling at the level of the entire set of binned reads. Specifically, a subset of reads of a given size was selected uniformly at random from the full collection of binned reads, and reconstruction and decoding were then carried out using only this subset.

## RESOURCE AVAILABILITY

### Lead contact

Requests for further information and resources should be directed to and will be fulfilled by the lead contact, Simonas Juzėnas.

### Materials availability

This study did not generate unique biological materials requiring distribution statements beyond the methods and supplementary information provided in this manuscript.

### Data and code availability

- Raw sequencing data generated in this study have been deposited in the European Nucleotide Archive (ENA) under accession number PRJEB88528.
- Analysis scripts and related code are available at https://github.com/genomika-lt/NinjaSeq.
- Any additional information required to reanalyze the data reported in this paper is available from the lead contact upon request.

## ACKNOWLEDGMENTS

The authors gratefully acknowledge financial support from the European Union through the European Innovation Council (EIC) Pathfinder Challenge: DNA-based digital data storage under the DNAMIC project (grant agreement No. 101115389) and the DiDAX project (grant agreement No. 101115134). Views and opinions expressed are however those of the authors only and do not necessarily reflect those of the European Union or the European Innovation Council Executive Agency. Neither the European Union nor the granting authority can be held responsible for them.

## AUTHOR CONTRIBUTIONS

Conceptualization: RP, LZ, IG, SJ; Data curation: KK, OS, HA, SJ; Formal analysis: SJ, OS, HA, TC; Funding acquisition: IG, ZY, LZ, SJ; Investigation: RP, KK, VG, IG, LZ; Methodology: RP, LZ, IG, SJ; Project administration: IG, EY, LZ, SJ; Resources: IG, EY, LZ; Supervision: LZ, EY, SJ; Visualization: SJ, OS, HA; Writing – original draft: SJ, RP, OS; Writing – review & editing: IG, OS, HA, KK, TC, VG, ZY, RP, LZ, EY, SJ

## DECLARATION OF GENERATIVE AI AND AI-ASSISTED TECHNOLOGIES

The authors declare that no generative AI or AI-assisted technologies were used to generate the scientific content or conclusions of this manuscript.

## SUPPLEMENTAL INFORMATION

**Supplementary Figure 1.**
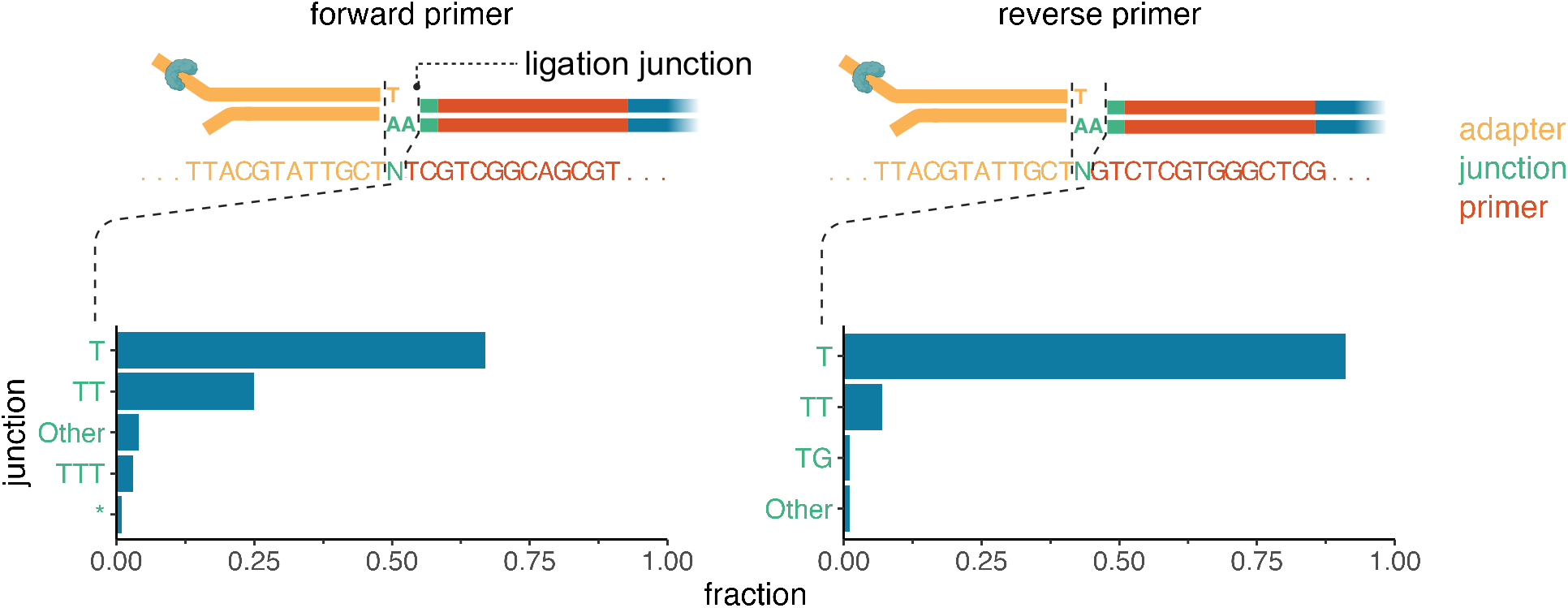
Sequence configurations at adapter-DNA junctions following BseGI restriction and ligation. Schematic representation of final sequence constructs obtained after BseGI digestion and adapter ligation. Yellow text indicates adapter sequence, red text denotes primer region sequences, and green text highlights the junction sequence between adapter and DNA following ligation. Left and right panels show the distribution of distinct ligation products generated using forward and reverse primers, respectively, illustrating the sequence heterogeneity resulting from the ligation process for each primer configuration.

**Supplementary Figure 2.**
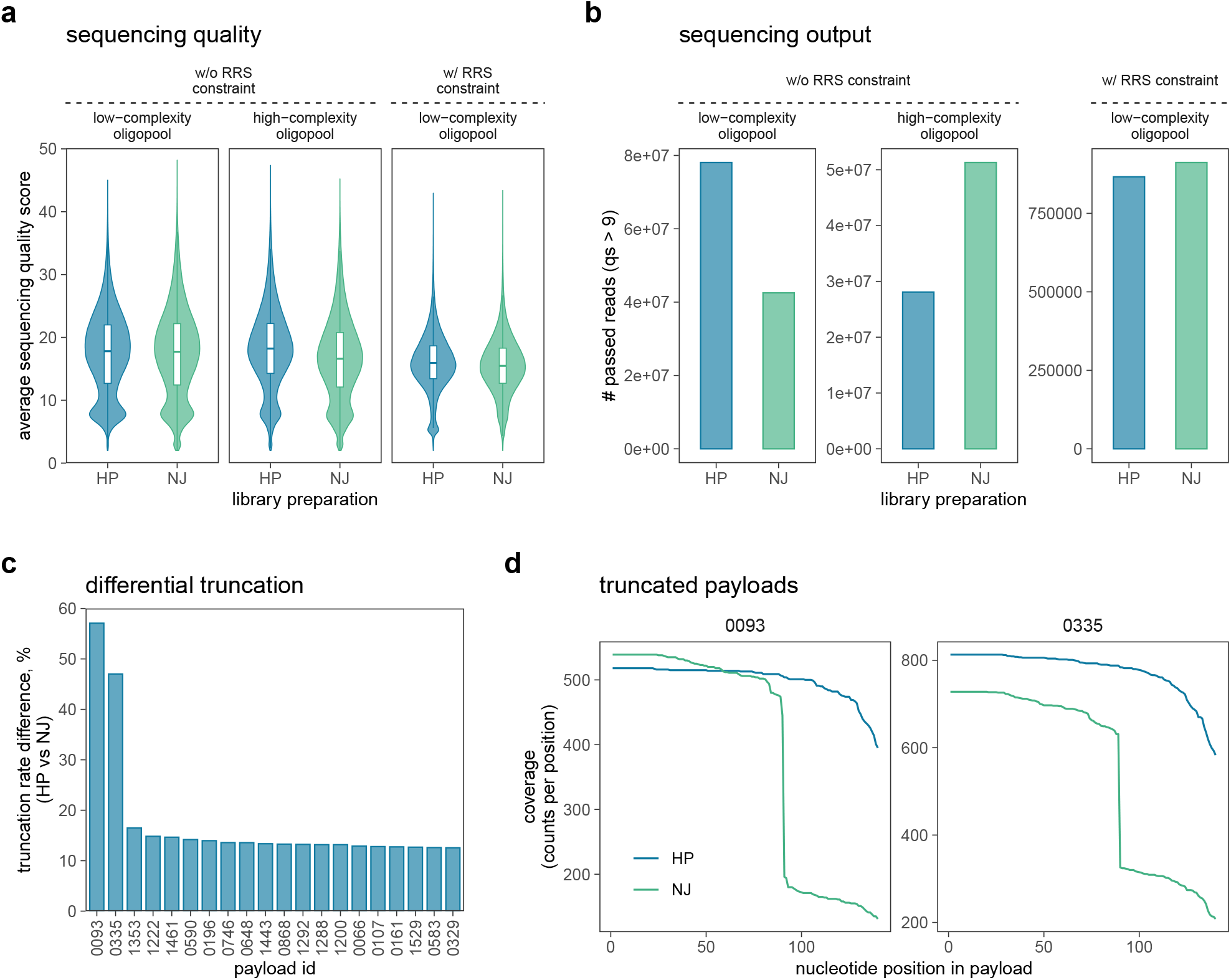
**a)** Sequencing quality score distributions. Violin plots show per read mean quality scores (qs) for HP and NJ libraries across oligopool types (low complexity or high complexity) and encoding designs (w/RRS constraint or w/o RRS constraint). Boxplots indicate the median and interquartile range. **b)** Sequencing output passing a quality threshold. Bar plots show the number of reads with qs*>*9 for each protocol, stratified by oligopool type and encoding design. **c)** Differential truncation across payloads in the RRS constrained pool. Bar plots show the top 20 payloads ranked by the absolute difference in truncation rate between protocols, reported as percentage points (HP vs NJ). **d** Coverage profiles for truncated payloads. Line plots show per position coverage (counts per position) along each selected payload for HP and NJ libraries; panels are faceted by payload identifier.

**Supplementary Table 1.**
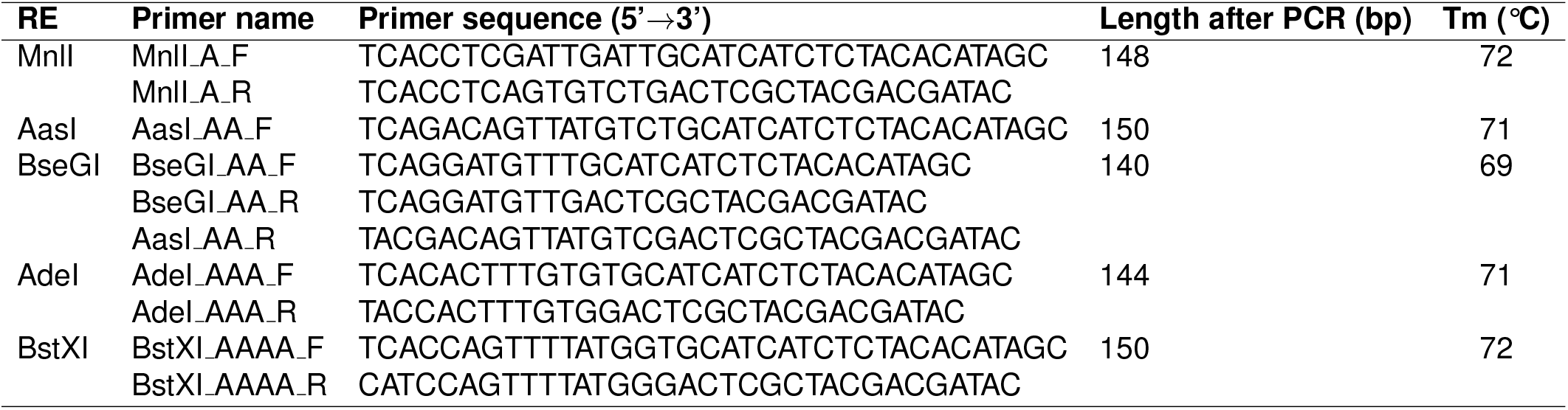
Primer sequences and PCR conditions for restriction enzyme experiments.

**Supplementary Table 2.** Restriction enzyme digestion conditions.

| Restriction enzyme | Reaction conditions | Thermal inactivation |
| --- | --- | --- |
| MnII | 37°C, 5 min | 65°C, 5 min |
| AasI | 37°C, 5 min | 80°C, 5 min |
| BseGI | 37°C, 5 min | 80°C, 5 min |
| Adel | 37°C, 5 min | 80°C, 5 min |
| BstXI | 37°C, 5 min | 80°C, 5 min |

**Supplementary Table 3.** Reagent cost comparison between the standard end-repair/dA-tailing workflow and the NinjaSeq workflow. Prices reflect manufacturer/distributor list prices in EUR (Lithuania); the sequencing kit and ligation module are common to both protocols. The end-preparation step alone is approximately 32-fold less expensive in NinjaSeq, contributing an overall per-reaction reduction of approximately 11% in reagent cost.

| Step | Reagent | Vendor | Price (EUR) | Reactions | Cost/rxn (EUR) |
| --- | --- | --- | --- | --- | --- |
| **Standard protocol** |  |  |  |  |  |
| End-preparation | NEBNext Ultra II End Repair/dA-Tailing Module | NEB | 391.0 | 24 | 16.3 |
| Ligation | NEBNext Quick Ligation Module | NEB | 411.0 | 20 | 20.6 |
| Sequencing | Ligation Sequencing Kit V14 (SQK-LSK114) | ONT | 650.0 | 6 | 108.3 |
|  |  |  |  | **Total** | **145.2** |
| **NinjaSeq protocol** |  |  |  |  |  |
| End-preparation | BseGI FastDigest restriction endonuclease | TFS | 51.0 | 100 | 0.5 |
| Ligation | NEBNext Quick Ligation Module | NEB | 411.0 | 20 | 20.6 |
| Sequencing | Ligation Sequencing Kit V14 (SQK-LSK114) | ONT | 650.0 | 6 | 108.3 |
|  |  |  |  | **Total** | **129.4** |
| **End-preparation step: 32-fold reduction. Total per-reaction reduction: 11%.** |  |  |  |  |  |

